# Kozak sequence libraries for characterizing transgenes across expression levels

**DOI:** 10.1101/2025.04.28.651141

**Authors:** Nidhi Shukla, Nisha D. Kamath, John C. Snell, Anna M. Bruchez, Kenneth A. Matreyek

**Affiliations:** Department of Pathology, Case Western Reserve University School of Medicine, Cleveland, Ohio, USA

## Abstract

Typical mammalian overexpression systems test protein sequence variants with little control over expression levels and steady-state protein abundances, hindering interpretations of how protein sequence and expression converge to yield phenotypic outcomes. We explored the translation initiation sequence, commonly referred to as the Kozak sequence, as a means to modulate protein steady-state abundance and cellular function. We performed sort-seq on a randomized library of the 6 nucleotides preceding the start codon, amounting to 4,042 sequences. Calibrating the scores revealed a ~100-fold range of protein steady-state abundances possible through manipulation of the Kozak sequence. We identified human germline variants with predicted expression-reducing Kozak substitutions in disease-associated genes. Modulating the cell surface abundance of the host cell receptor ACE2 controlled the rate at which those cells became infected by SARS-like coronavirus spike pseudotyped particles. We demonstrated the potential of the approach by simultaneously testing Kozak libraries with a small panel of coding variants for ACE2 and STIM1. This approach lays the methodological groundwork for linking the causal relationships between protein sequence, abundance, and functional outcome.

## Introduction

Advancements in recombinant DNA technologies have aided development of cell-based assays characterizing how expression of transgenic coding sequences can impact cellular processes of interest. Unfortunately, most genetic experiments are performed in a binary fashion by simply toggling the existence or absence of a protein without major consideration toward controlling the amount of protein product being created per cell. We lack the experimental tools and knowledge needed to dial-in the precise levels of protein abundance and quantitatively observe the phenotypic outcomes.

For synthetically engineered systems, the Kozak sequence, also known as the eukaryotic translation initiation sequence (TIS), presents a convenient, genetically encoded method for modulating the steady-state abundance of a transgenic protein of interest. The Kozak sequence dictates the efficiency of cap-dependent protein translation by controlling the likelihood that a ribosome initiates at a particular ATG start codon encountered upon scanning from the 5’ end of the mRNA (Kozak, 1981, 1989). The Kozak sequence is generally regarded as the 6 nucleotides preceding the start codon, designated the −6 to −1 positions, as well as the first nucleotide following the start codon, designated the +4 position (Kozak, 1987), with other nearby nucleotides potentially contributing minor effects (Noderer *et al*, 2014).

Various studies have measured relative differences in efficiency between Kozak sequences across contexts. The first large-scale eukaryotic translation initiation dataset for Kozak sequences was created in PD-31 mouse pre-B lymphocytes, yielding a position weight matrix model for predicting translational score (Noderer *et al*, 2014). These scores were pseudo-quantitative, as the fold difference in protein amounts that could be inferred by comparing two scores was never defined. Other studies experimentally tested dozens to hundreds of Kozak sequences in yeast (Petersen *et al*, 2018), Drosophila Kc167 cells (Acevedo *et al*, 2018), and human HEK 293T cells (Ambrosini *et al*, 2022). Despite these studies demonstrating the relative universality of Kozak sequence effects across eukaryotic cell backgrounds, we still lack a quantitative understanding for how any given pair of Kozak sequences differ in their resulting steady-state protein abundances.

We addressed this by creating a library of ~4,042 Kozak sequences modulating the expression level of the cell-surface protein ACE2 with the fluorescent protein miRFP670 fused to its cytoplasmic tail. These sequences were functionally assessed and given a calibrated abundance score, reflecting the mean-fluorescence intensity emitted by miRFP670 translated from that Kozak sequence. We used the ensuing scores to identify human germline variants that are potentially pathogenic due to reduced total amounts of particular disease associated protein. We varied ACE2 expression and tested the cells for their susceptibility to infection by virus particles dependent on ACE2 for entry into the cell. We also varied the expression amounts of STIM1, a protein involved in calcium signaling and cellular metabolism, and evaluated its effects on cell survival. This work demonstrates the potential of Kozak sequence modulation for learning how varying expression levels of different transgenic sequences or sequence variants correspond to functional output.

## Results

### Single-copy integration of a fluorescent protein Kozak library

To perform our transgenic manipulations in cultured human-derived HEK293T cells, we harnessed targeted single-copy plasmid integration for transgenic overexpression, using a Bxb1 landing pad cassette (Matreyek *et al*, 2017, 2020; Shukla *et al*, 2021). This system relies upon cells with a prior genomic integration of a single Bxb1 recombination site. The site is preceded by a Tet-inducible promoter, and followed by a cassette encoding a blue fluorescent reporter, positive and negative selectable markers, and the Bxb1 recombinase enzyme (**Fig 1A, top**). Expansion of clonal “landing pad” cells yields an isogenic line usable for integrating transgenic elements of interest.

**Figure 1.**
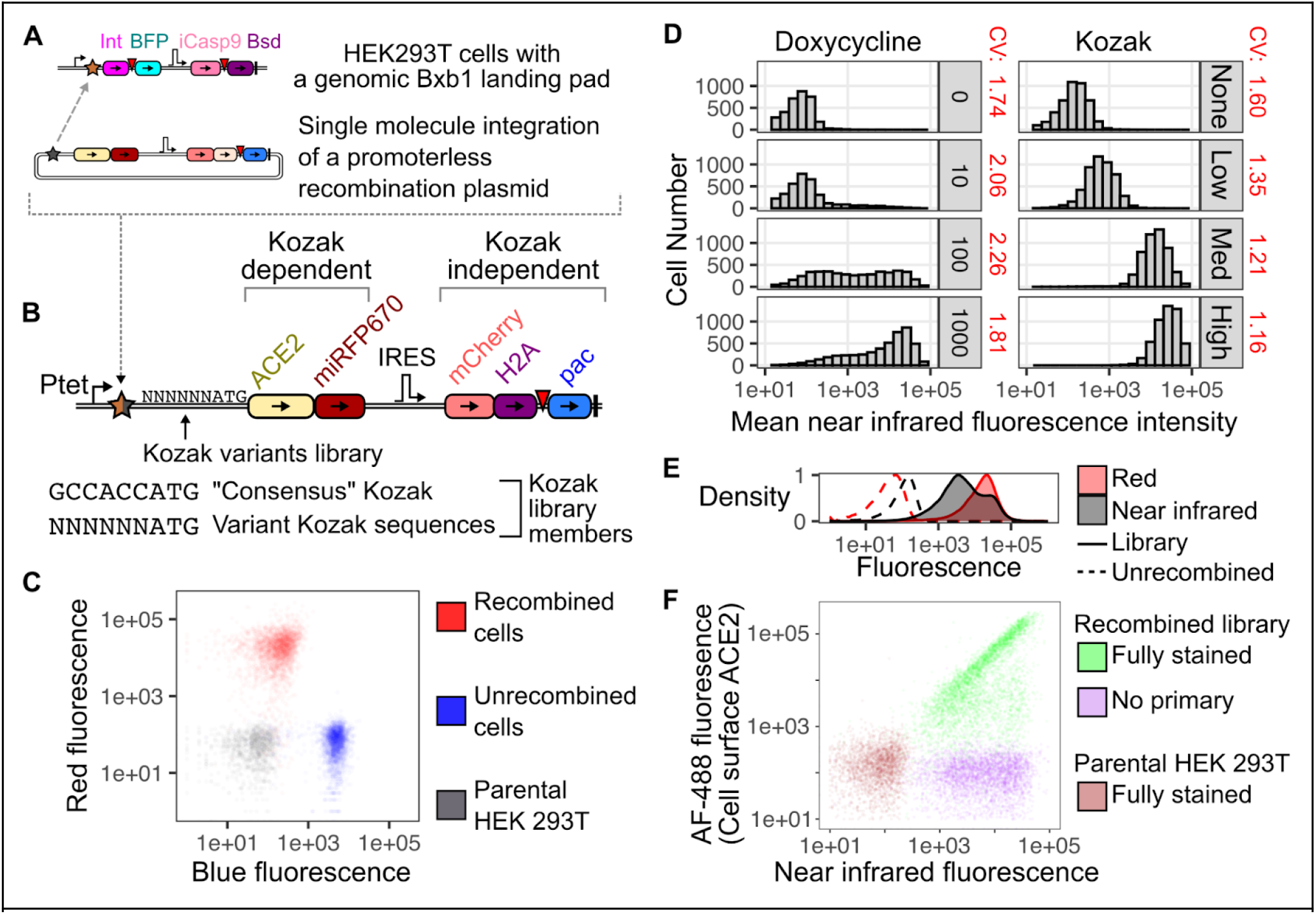
Single copy integration of a Kozak variant library. A) Schematic of the HEK 293T landing pad system, with a single plasmid molecule (bottom) integrating into a single genomically engineered site (top). B) Schematic of the transgenic DNA present following plasmid integration. A degenerate Kozak sequence library precedes ACE2 and miRFP670. A histone-fused mCherry and puromycin resistance gene (pac) is encoded behind an IRES and are translated independent of the Kozak sequence. Ptet, Tet inducible promoter; IRES, internal ribosome entry site; upside down red triangle, 2A translation stop-start sequence; brown and black stars, Bxb1 attP and attB recombination sequences, respectively. C) Representative blue and red fluorescence flow cytometry profiles for unmodified HEK 293T cells (black), unrecombined landing pad cells (blue), and antibiotic selected recombined cells (red). D) Representative near infrared fluorescence from high miRFP670 expressing cells treated with a titration of doxycycline inducer (left), or cells expressing miRFP behind Kozak sequences of various translational strengths (right). E) Cell surface ACE2 protein staining of cells expressing the ACE2-Kozak library (green) or control conditions lacking the primary antibody (purple), and unmodified HEK 293T cells mixed with both primary and secondary antibodies (brown).

Transfection of these cells with promoterless plasmids encoding the complementary Bxb1 recombination site inserts a single plasmid molecule into the existing genomic landing pad site (**Fig 1A, bottom**). The previously promoterless transgene of interest is now behind the Tet-inducible promoter (**Fig 1B)**, displacing and turning off expression of the prior DNA cassette. The recombined cells encoding the transgene of interest can be identified due to their loss of blue fluorescence and simultaneous gain of red fluorescence (**Fig 1C**). This system allows highly precise and stable transgenic expression compared to lentiviral transduction and transient transfection methods (Shukla *et al*, 2021). Since only a single molecule from a large pool of transfected plasmids becomes integrated, this transgenic expression method is compatible with single-cell library methods requiring strict links between cell genotype and phenotype (Matreyek *et al*, 2017, 2018, 2020).

Controlled modulation of steady-state transgenic protein expression is key to quantitative understanding of protein function, but generalizable strategies remain underdeveloped. Traditional approaches typically harness inducible promoters, such as the Tet-inducible promoter, where various concentrations of the chemical inducer doxycycline is used to achieve intermediate amounts of average protein expression at the population level. These methods are imprecise, as there is large cellular heterogeneity in protein abundance with intermediate inducer amounts (**Fig 1D, left**). The observed range of protein products achieved with intermediate chemical inducer concentrations are also not genetically encoded and cannot be measured using scalable molecular readouts such as high throughput DNA sequencing.

To achieve controlled modulation of protein steady-state abundance for our gene of interest, we focused on alterations to the eukaryotic translation initiation sequence. The Kozak sequence determines the rate of cap-dependent protein translation for a protein from a given mRNA, corresponding to a specific steady-state protein abundance setpoint (Noderer *et al*, 2014). We confirmed that altering the Kozak sequence could tune steady-state protein abundance, and was substantially more homogenous than with doxycycline titration, with more consistent coefficients of variation across the expression range (**Fig 1D, right**).

As each Kozak-dictated steady-state abundance phenotype is genetically encoded, we used DNA sequencing to identify cells with different levels of transgenic protein. To test the full range of protein abundances within a single experiment, we created a library of Kozak sequence variants using a primer with an internal degenerate segment encoding all possible nucleotides at the 6 positions preceding the start codon. As alteration of the +4 position changes the protein coding sequence, we kept this nucleotide invariant in our experiments, opting to maintain the wildtype nucleotide encoded by the open reading frame. This library had a total potential complexity of 4^6^ or 4,096 members (**Fig 1B, bottom**).

To experimentally test the Kozak sequences, we created a promoterless recombination plasmid encoding the cell-surface protein ACE2 with the near-infrared fluorescent protein miRFP670 fused to its cytoplasmic C-terminus (**Fig 1B, top**). The coding region was immediately preceded by the degenerate Kozak library nucleotides, which determined the translation rate and steady-state abundance of the ACE2-miRFP670 fusion protein. To mark the genetically modified cells independently of the Kozak-dependent translation products, the recombinant DNA construct also encoded the red fluorescent protein mCherry fused to histone 2A, cotranslationally linked to the puromycin resistance gene. These coding regions were preceded by an internal ribosome entry site, and thus translated independently from the Kozak sequence (**Fig 1B, top**). The plasmid library was recombined into the HEK 293T landing pad cells and enriched using sequential negative and positive selection to yield a pure population of successfully modified, uniformly red fluorescent cells (**Fig 1E**). In contrast, these cells had a wider range of near infrared fluorescence due to differing rates of cap-dependent translation from the Kozak sequence library (**Fig 1E**).

We confirmed that miRFP670 intensity reflected the amount of ACE2 present on the cell surface by staining the cells with an ACE2-targeting antibody followed by an Alexa Fluor-488-conjugated secondary antibody. We observed a clear linear relationship between near infrared fluorescence from miRFP670 and green fluorescence from antibody staining (**Fig 1F**). This signal was specific for overexpressed ACE2, as omission of either the primary or secondary antibody ablated green fluorescence, and unmodified parental HEK 293T cells did not stain with the full combination of primary and secondary antibody. We proceeded by using near infrared fluorescence intensity emitted by miRFP670 to minimize unnecessary cell manipulation and costs during subsequent assay steps.

### Quantitating Kozak sequence impact on protein abundance

We confirmed the range of protein abundances possible with the Kozak sequence library by recombining plasmids encoding known strong and weak Kozak variants as controls (Noderer *et al*, 2014). The strong consensus Kozak (GCCACCATG) drove high miRFP670 fluorescence, while a weak Kozak (TTATGGATG) yielded low miRFP670 fluorescence (**Fig 2A**). The cells within the library population spanned these levels of fluorescence (**Fig 2A, green**), consistent with the library encoding a diverse range of translational efficiencies possible with this approach.

**Figure 2.**
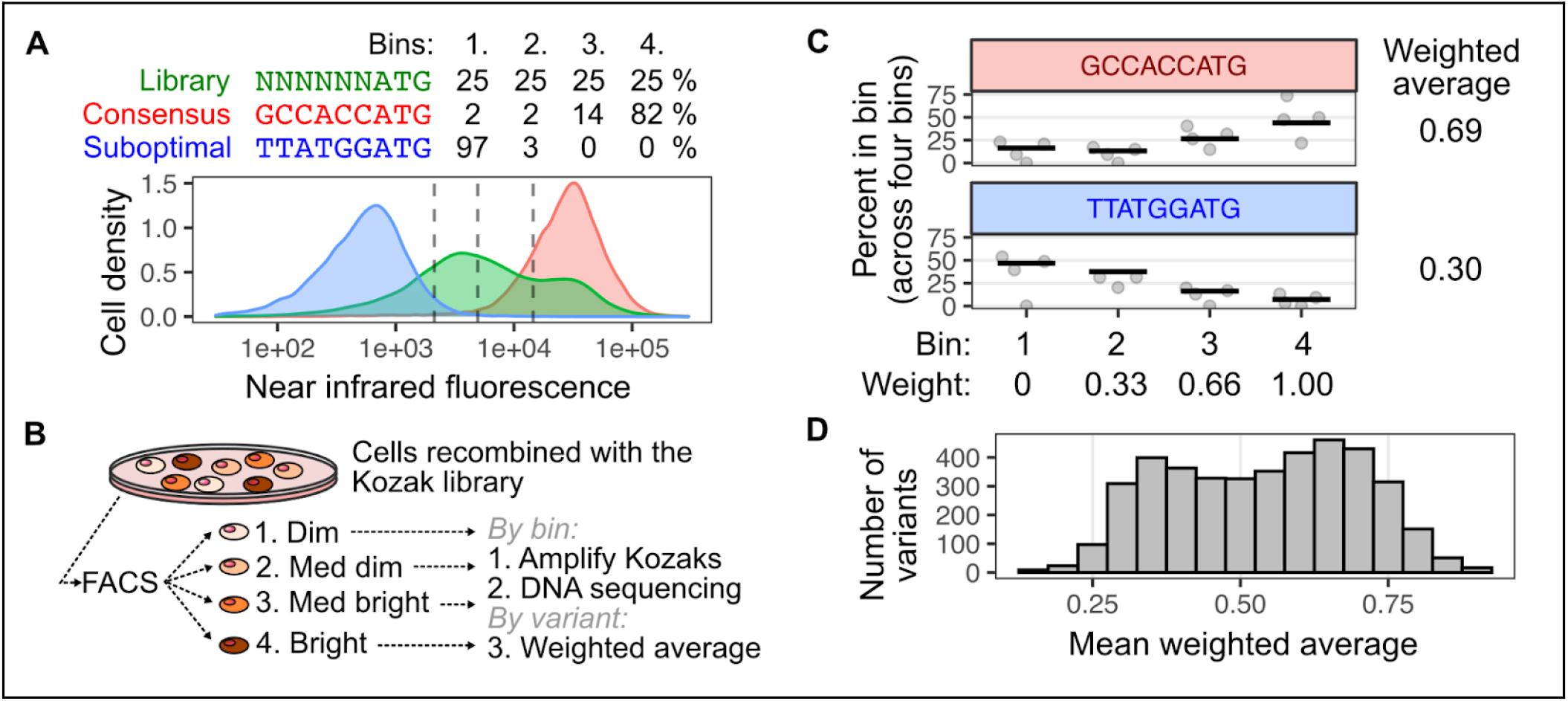
4-way sorting and sort-seq of the ACE2-Kozak cells based on near infrared fluorescence. A) Cells encoding the ACE2-Kozak library (green) were compared to cells encoding ACE2 behind a weak (blue) or strong (red) translation Kozak sequences, and assessed for near infrared miRFP670 fluorescence using flow cytometry. The ACE2-Kozak library cell population was broken into quartile bins using threshold values (dotted lines), and the distribution of the control cells amongst the bins are shown as percentages. B) Schematic of the Sort-seq approach used to calculate steady-state abundance values corresponding to each Kozak variant. C) The distribution of DNA sequencing reads corresponding to the control Kozak sequences amongst the four bins for 5 replicate experiments, as well as the calculated mean weighted average corresponding to each sequence. D) Histogram of the mean weighted averages for all Kozak variants in the dataset.

To characterize the steady-state protein abundance exhibited by each member of the library, the population of expanded library cells were separated into equally-populated quartiles using fluorescence activated cells sorting (FACS), creating four subsets of cells with different levels of fluorescence (**Fig 2A, 2B**). The distribution of the cells across the four bins reflected the translation rate exhibited by that Kozak variant. For example, 97% of the TTATGGATG control Kozak cells were in the lowest quartile bin, with the remaining 3% in the second lowest bin, whereas 82% of the GCCACCATG control Kozak-containing cells were in the highest bin, with 14% found in the second highest bin, and the remaining 4% equally distributed between the lowest two bins (**Fig 2B**). The library was independently sorted into quartile bins on 4 separate occasions.

We performed high throughput DNA sequencing to quantitate the distributions of the library members (**Figure 2B**). Upon sorting, the cells from each quartile bin were grown for a week after which their genomic DNA was extracted. The DNA encompassing the Kozak sequence was amplified by polymerase chain reaction (PCR), and sequenced with Illumina high throughput DNA sequencing. The read counts for each Kozak sequence variant in each bin was divided by the total number of reads in that bin to convert each Kozak sequence variant into a frequency. By multiplying the frequencies with a weight value specific for each bin (ranging from 0 to 1; see **Fig 2C**), each Kozak sequence variant was given a weighted average value.

This process was repeated for each sorted replicate, and a mean weighted average was calculated for each variant across the replicate experiments. We obtained weighted average scores for 4,042 variants. The variants largely spanned the range from roughly 0.13 to 0.91 (**Fig 2D**). The aforementioned strong and weak Kozak control variants exhibited weighted averages of 0.69 and 0.30, respectively (**Fig 2C**).

While these weighted averages can distinguish the effects conferred by different Kozak sequences, they cannot be directly used to interpret fold differences in protein steady-state abundances. We thus calibrated our weighted average values to the mean fluorescence intensity (MFI) exhibited by miRFP670 when tested individually. We chose a panel of 16 Kozak sequences spanning the range of weighted scores, created constructs specific for each sequence, generated clonal populations of cells with each construct, and assessed the resulting cells for miRFP670 fluorescence using flow cytometry. The resulting MFIs exhibited high correlation with the mean weighted averages (Pearson’s r^2^ = 0.92, Spearman’s ρ^2^ = 0.86; **Fig 3A**). We used regression to calculate an estimated MFI for each variant scored in the library, and refer to these values as “calibrated abundance scores”.

**Figure 3.**
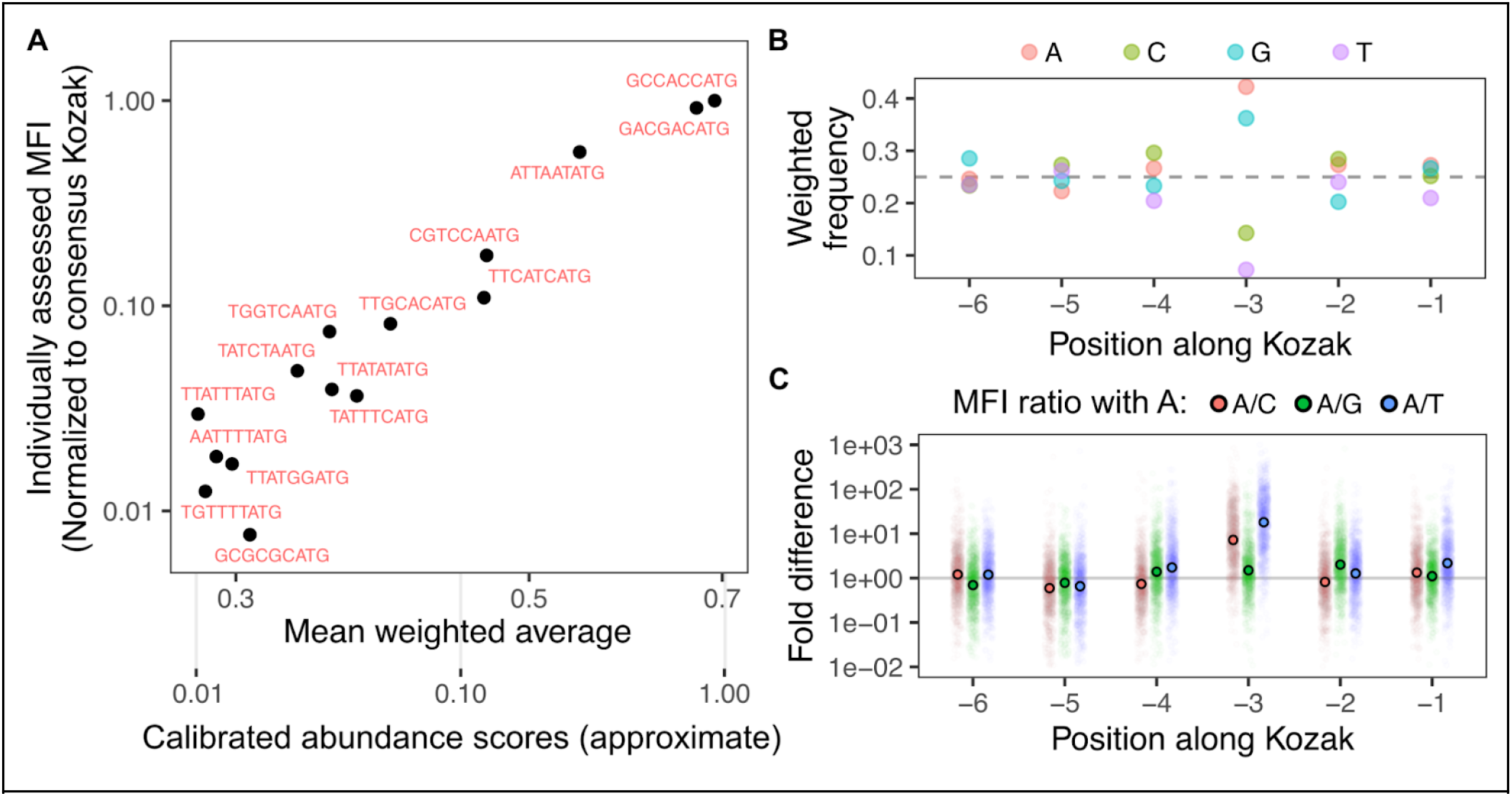
Calibration and analysis of Kozak sort-seq scores. A) Comparison of the mean weighted average of each Kozak sequence (x-axis, top) from sort-seq analysis, as well as the individually assessed mean fluorescence intensity of cells encoding each Kozak through analytical flow cytometry (y-axis), normalized to the GCCACC consensus Kozak sequence. Based on this calibration curve, an estimated MFI was calculated based on the sort-seq values (x-axis, bottom). B) Position weight matrix denoting individual nucleotide contributions to overall Kozak sequence effects on calibrated abundance score. C) Positional impacts of various nucleotides on translational efficiency, shown as fold differences compared to that from A at each position. Small colored circles denote individual variants from the library, while the larger circles with black outlines denote geometric mean values.

These values enabled the generation of new quantitative magnitude estimates for known positional effects within Kozak sequences. As expected, the position weight matrix values showed that adenine and guanine were highly favored at position −3 (**Fig 3B**). The importance of this position is well known (Noderer *et al*, 2014; Kozak, 1986, 1987), but our MFI-scaled dataset yielded alternative ways to measure this effect. For every sequence, we calculated the fold difference in MFI-calibrated score when A was encoded at a given position, compared to C, G, and T (**Fig 3C**). On average, we found that an A at the −3 position yielded 1.3-fold more protein abundance over G, 5.2-fold over C, and 13.4-fold over T. In contrast, the effects at position −1 were comparatively minor, with an A yielding 1.1-fold more protein abundance over G, 1.3-fold over C, and 2.1-fold over T.

### Comparison of scores across data sources

We compared our dataset of calibrated abundance scores to the Noderer *et al.* dataset. Their reported scores were outputs of a position weight matrix model based on their experimentally measured data (Noderer *et al*, 2014). Their modeled scores correlated with our calibrated abundance scores (Pearson’s r^2^ = 0.56, Spearman’s ρ^2^ = 0.6; **Fig 4A**), and the correlation improved when the logarithm of our calibrated abundance scores were used (Pearson’s r^2^ = 0.67). Thus, their scores, ranging from 11 to 161, were logarithmically proportional to more traditional measurements of relative protein abundance such as with flow cytometry.

**Figure 4.**
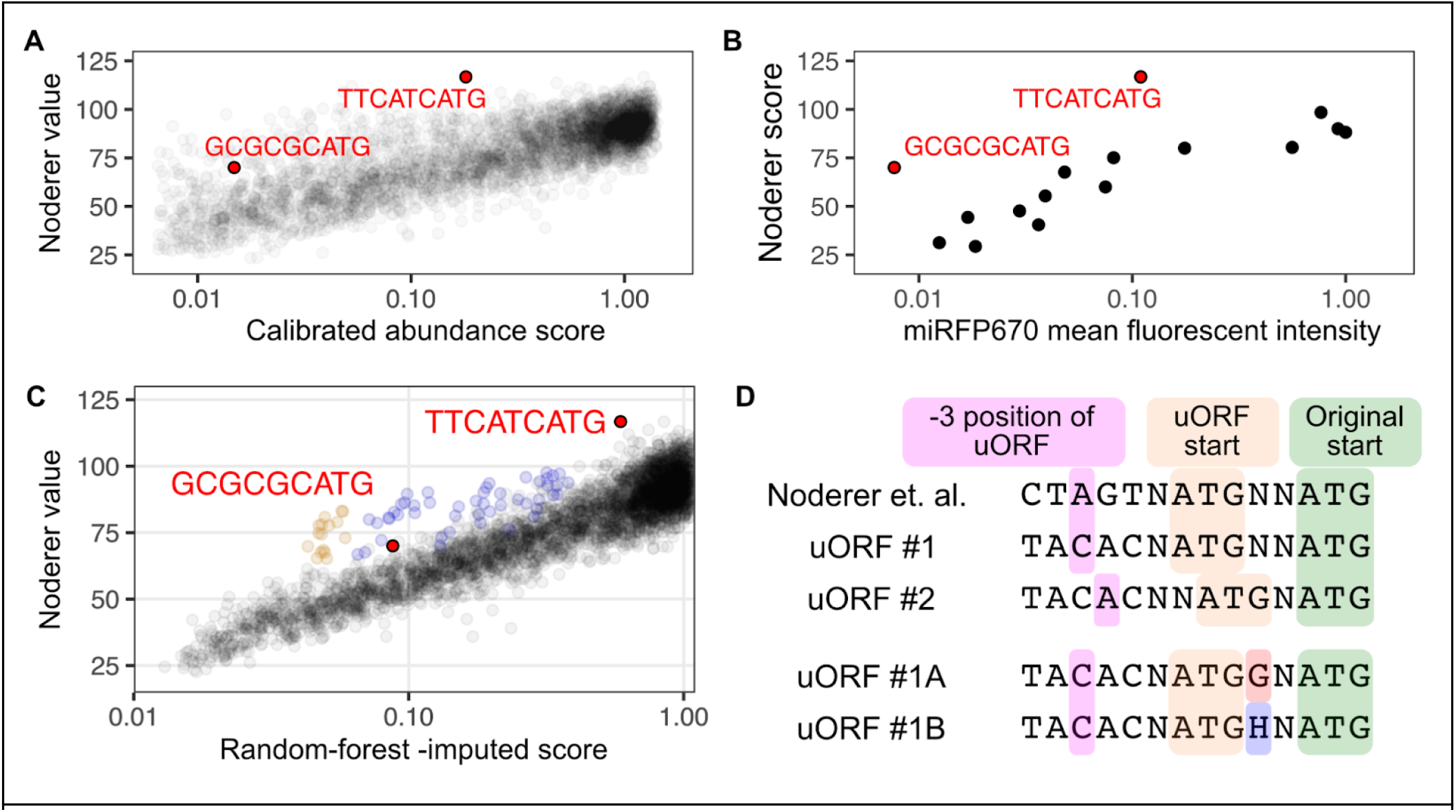
Comparison to Kozak sequences tested in other model systems. A) Comparison of our calibrated abundance scores with mean Kozak translational scores from the Noderer et al. dataset. B) Geometric means of miRFP670 mean fluorescence intensity measured when the indicated Kozak sequences were tested individually, and comparison with the Noderer et al. dataset. C) Comparison of the random-forest-imputed abundance scores with the Noderer et al translational efficiency scores. Two subtypes of Kozak sequences encoding upstream open reading frames starting at the −5 position, both defying the correlative pattern of the overall dataset, are colored orange and blue. D) Schematic showing the nucleotide sequences upstream of the degenerate Kozak library site, demonstrating the sequence features likely underlying reasons for discordance between the two datasets. uORF#1A and uORF#1B corresponds to the orange and blue points in panel C, respectively.

Despite the overall correlation, there were some discrepancies between the datasets. This was particularly evident for “GCGCGCATG” and “TTCATCATG, as these were also present in our panel of individually tested sequences (**Fig 4B**). We suspected that the Noderer *et al.* scores being reported as position weight matrix output values may have contributed to the discrepancies between our two datasets. We created a similar set of modeled output values using a random forest machine learning model on the MFI-calibrated scores yielded by our assay. We observed model outputs that were highly correlated with the Noderer *et al* values (Pearson’s r^2^ = 0.85, Spearman’s ρ^2^ = 0.75). The improved correlation of the random forest model output, as compared to our calibrated scores, suggest that the models converged on similar Kozak sequence properties.

A subset of data points were outliers exhibiting higher Noderer scores than predicted by our model (**Fig 4C, orange and blue points**). The biggest discrepancies were from sequences introducing an ATG start codon prior to the intended protein start site, creating out-of-frame upstream open reading frames (uORFs) at the −5 position, with those encoding G at the +4 position the most extreme (**Fig 4C**). A G at the +4 position is predicted to enhance translation initiation of the uORF (Noderer *et al*, 2014), which likely causes greater disruption of the downstream coding ORF than other nucleotides at the +4 position. The discrepancy between the models is likely attributable to there being a weaker G nucleotide in the −3 position of the Noderer et al. constructs, while our constructs encode the translation-enhancing A nucleotide (**Fig 4D**). Thus, these scores can also inform how competition with uORFs impact the steady-state abundance of transgenically encoded proteins.

We assessed the generality of these effects by comparing our scores to other studies testing Kozak sequence changes at smaller throughput across model systems. Our scores correlated with 244 values derived from lentivirally transduced HEK 293T cells (Pearson’s r^2^: 0.55, Spearman’s ρ^2^: 0.56) (Ambrosini *et al*, 2022) (**Supplementary Fig 1**). Our scores also correlated with a set of 50 constructs tested in insect cells (Pearson’s r^2^: 0.74, Spearman’s ρ^2^: 0.77) (Acevedo *et al*, 2018). Thus, our scores are likely to be applicable when modulating Kozak sequences across eukaryotic cells.

### Germline Kozak variants in disease genes

We applied our calibrated scores to identify human germline variants that may cause disease through reduced steady-state abundance of key proteins. We identified Kozak sequence variants reported in ClinVar, a repository of largely germline genetic variations and their clinical significance observed through clinical laboratory genetic testing of patients (Landrum *et al>*, 2018). We found 1,826 Kozak variants in ClinVar that were within the 6 nucleotide stretch scored within our dataset. Roughly 5% of these were predicted to reduce protein abundance by at least 4-fold (**Fig 5A**).

**Figure 5.**
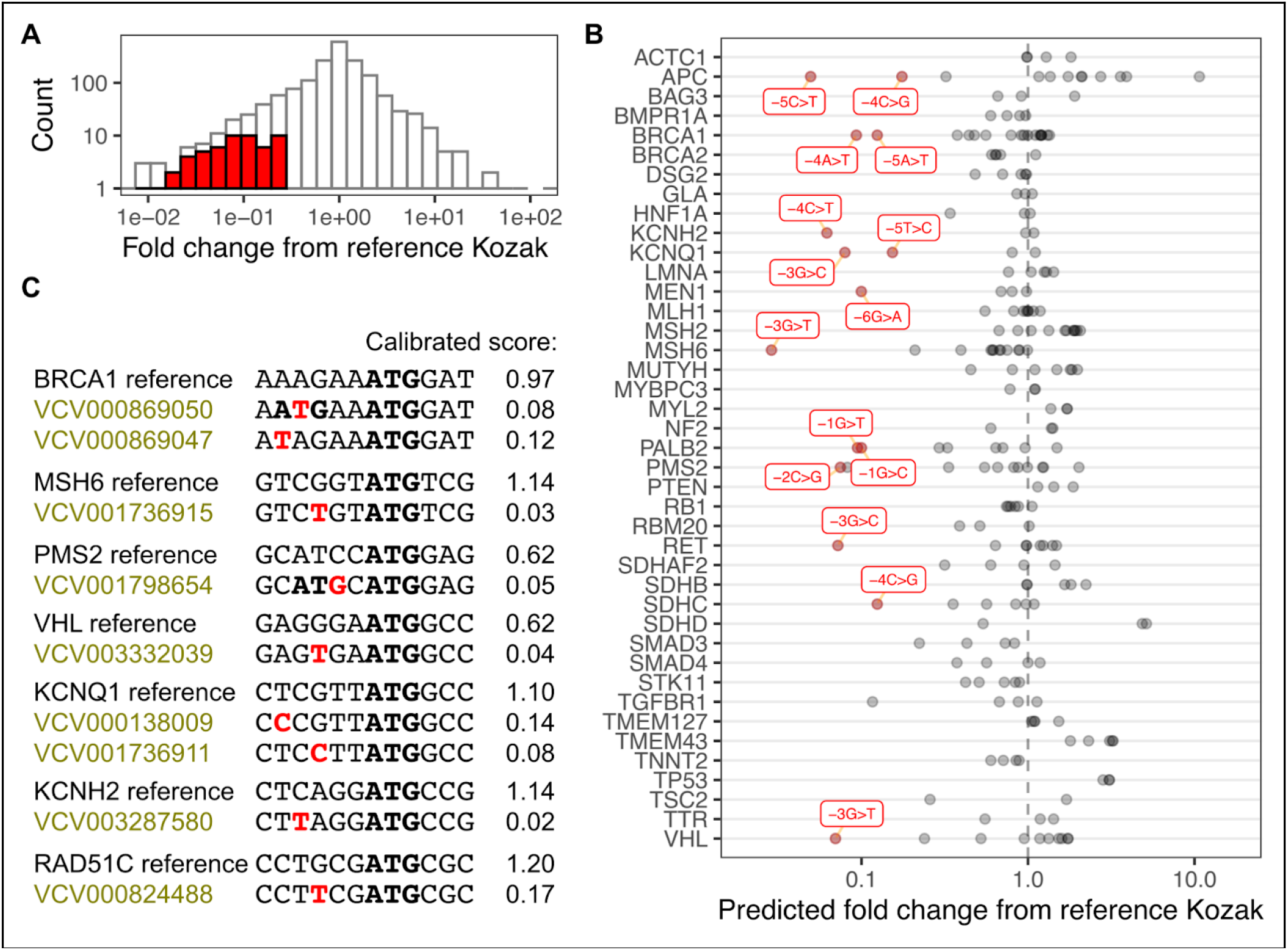
Germline Kozak variants in disease genes. A) A histogram of Kozak sequence variants listed in ClinVar, with high confidence scores causing greater than 4-fold decreases to calibrated abundance score relative to the reference sequence colored as red bars. B) Plot denoting protein relative fold abundance changes predicted for various ClinVar entries of Kozak germline variants, for genes found in the ACMG73. High confidence scores exhibiting greater than 5-fold reduction to calibrated abundance score from the reference sequence are labeled and colored in red. C) Examples of various disease-associated genes, their reference Kozak sequences, and single nucleotide variants observed as entries in ClinVar. The single nucleotide variant is colored red, and initiator methionine codons are in bold. Labels for variants of uncertain significance are colored gold.

We focused on the ACMG73, which is a collection of genes most important for clinical genetics (Miller *et al*, 2023). We found 18 Kozak variants in these genes expected to reduce protein abundance by at least 5-fold (**Fig 5B; red points**). Kozak variants in tumor suppressor genes were particularly notable, as the presence of even one variant allele encoding a protein with reduced activity can result in increased cancer predisposition. This included multiple DNA repair genes listed in the ACMG73, such as BRCA1, MSH6, PMS2, and TP53 (**Fig 5B**).

The variants with the greatest magnitude reductions generally encoded uORFs, such as the −4A>T variant of BRCA1 and the −2C>G variant of PMS2 (**Fig 5C**). The functional impact of this PMS2 variant was recently shown to exhibit significantly reduced translation in an alternative assay system (Matoy *et al*, 2024). Impactful substitutions of the −3 position were also observed, such as −3G>T variants of MSH6 and VHL, and a −3A>T variant of KCNH2. There were also variants in established disease genes not included in the ACMG73, such as RAD51C. As these variants are currently listed as “variants of unknown significance”, the functional results from our dataset may be helpful in future reclassification of these types of variants (Starita *et al*, 2017).

### Gradations of functional output with Kozak libraries

With the relative steady-state protein abundances defined for each Kozak sequence, we used the Kozak sequence library in cultured cell assays to characterize how different protein abundance levels impact cellular activity. Using a handful of Kozak sequences, we previously showed that the cell surface abundance level of ACE2 can impact virus entry and rates of antibody-mediated neutralization of SARS-CoV-2 spike pseudotyped virus particles (Shukla *et al*, 2021; Farrell *et al*, 2022; Roelle *et al*, 2022). We used the Kozak library driving ACE2-miRFP670 expression to assess how virus infectivity changes across the full range of expression levels.

To assess how ACE2 cell surface abundance correlated with infectivity, we performed a multiplexed infection assay with our ACE2 Kozak library cells (Shukla *et al*, 2024) (**Fig 6A**). We mixed the cells with lentiviral particles encoding EGFP and the hygromycin resistance gene, but coated with SARS-CoV spike, SARS-CoV-2 spike, or the unrelated glycoprotein from vesicular stomatitis virus (VSV-G). Unlike the coronavirus spike proteins, VSV-G uses the ubiquitously expressed low density lipoprotein receptor for cell entry rather than ACE2 (**Fig 6A**). After green cells were observed, genomic DNA from a subsample of cells were collected and stored. The remaining cells were treated with hygromycin to remove uninfected cells, and ensure that only genomic DNA from infected cells would be retained. The Kozak sequences were amplified, sequenced, and tallied before and after selection. An enrichment ratio was calculated for each Kozak sequence to determine their relative rates of infection by each pseudotyped virus (**Fig 6A**).

**Figure 6.**
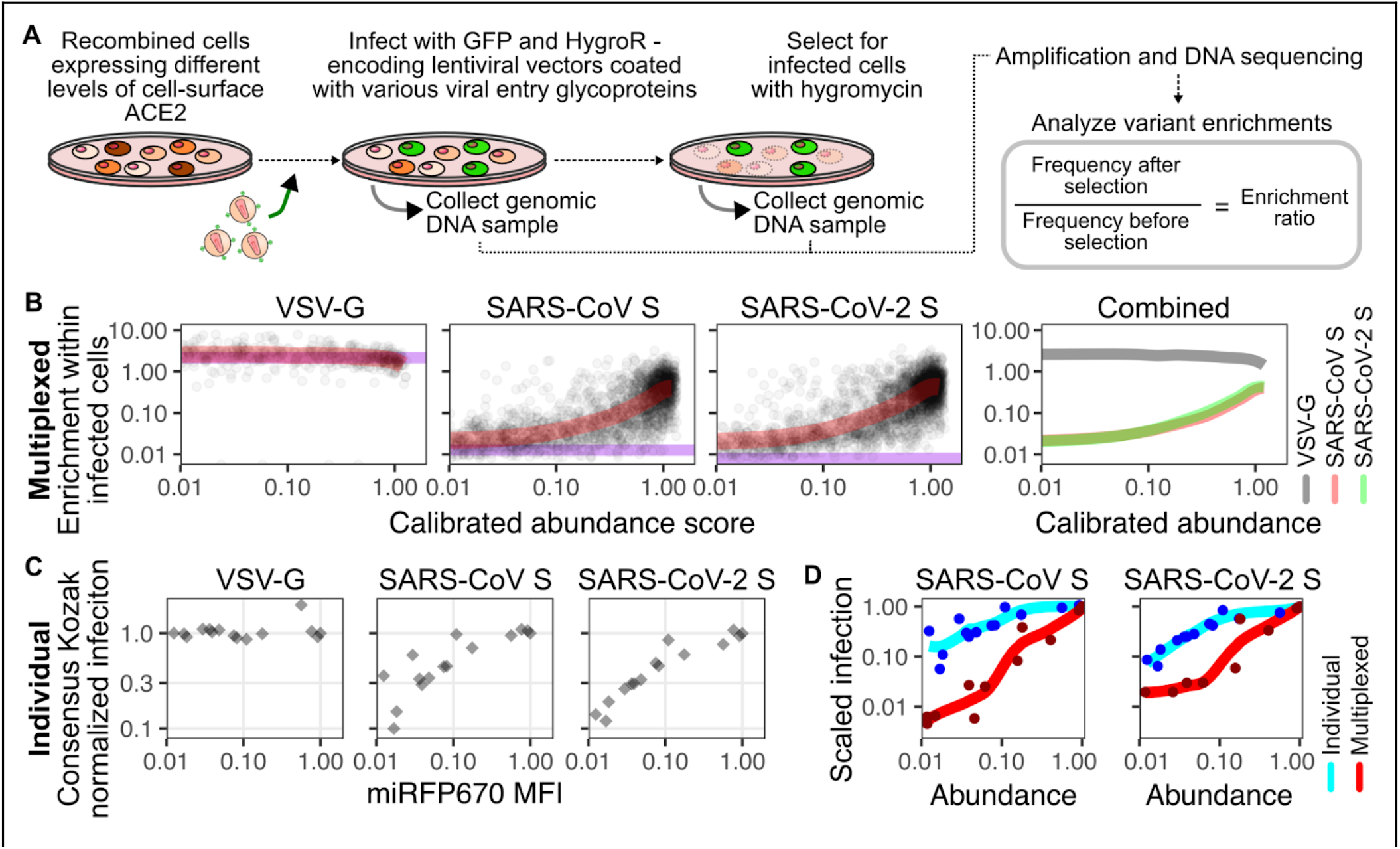
ACE2 expression level impacts on SARS-CoV spike-mediated infection. A) Work-flow diagram describing key physical and computational steps for the multiplex infection assay readout. B) Scatterplots of pseudotyped virus infectivity as a function of calibrated abundance scores. The semi-translucent red lines denote a LOESS fit to the points. The plot to the rightmost column overlay these LOESS curves. The horizontal purple line denotes the infectivity of template plasmid, which encodes a null ACE2 sequence lacking its start codon. C) Scatterplots of pseudotyped virus infectivity as a function of miRFP670 fluorescence of a set of clonally tested Kozak sequences. D) Scatterplot overlays of the same set of Kozak sequences shown in panels 6B and 6C, with normalized infection score as a function of protein abundance. The LOESS fit colors denote the two different assay types compared.

Visualizing the enrichment ratio relative to our previously ascertained abundance values revealed how ACE2 cell surface amounts corresponded to infection (**Fig 6B**). As expected, pseudotyped viruses coated with VSV-G infected all cells equally, regardless of the level of ACE2 abundance (**Fig 6B**). In contrast, infectivity of the SARS-CoV and SARS-CoV-2 spike pseudotyped viruses correlated with the level of ACE2 abundance, and were nearly identical when compared to each other (**Fig 6B, right**).

We next compared whether the relationship between ACE2 abundance and likelihood of pseudotyped virus entry changed depending on the assay format. We thus infected the same 15 clonal cell lines of varying Kozak strengths used to create the MFI calibration scores with the pseudotyped viruses (**Fig 6C**). When the results from each assay format were scaled to the consensus Kozak, we observed clear differences imposed by ACE2 expression on target cell infection, depending on whether the assay was performed with homogeneous or heterogeneous cell mixtures (**Fig 6D**). For example, ACE2 Kozak sequences that had calibrated abundance scores of 0.1 exhibited 10-fold less infection than ACE2 behind a consensus Kozak when tested as a pooled library, while the same sequences when tested as individual, homogenous cultures exhibited no differences to infectivity (**Fig 6D**). These results highlight discrepancies in biological assays that can only be identified when performing the assay in a pooled manner.

### Modulating protein sequence and abundance

Simultaneous modulation of protein sequence and steady-state abundance levels would vastly improve our understanding of the functional impacts of coding sequence changes. Our Kozak library experiments showed that the −6 position has little impact on determining ensuing protein steady-state abundance, so we explored rendering this nucleotide invariant to serve as a genetically encoded identifier for marking which construct variant is being sequenced, like a standard nucleotide barcode. We found that all of the remaining library members still spanned the full 100-fold range of abundance, regardless of which nucleotide base was fixed at the −6 position (**Fig 7A**).

**Figure 7.**
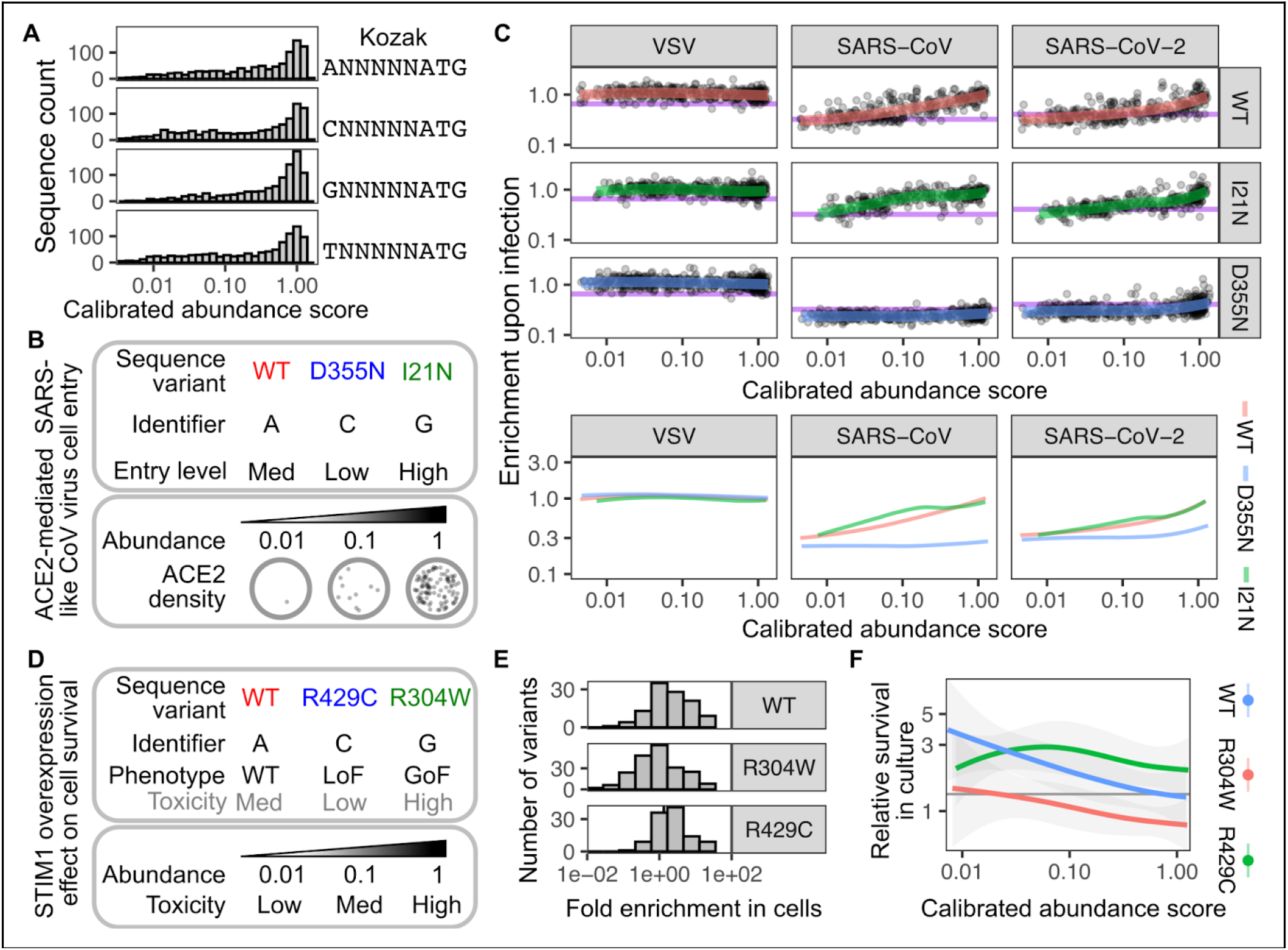
Results of simultaneously modulating protein sequence and expression. A) Histograms of calibrated scores when the −6 Kozak nucleotide is left invariant. B) Schematic demonstrating the ACE2 sequence variants, their nucleotide identifiers, and their anticipated infection phenotypes (top), as well as a visual depiction of 100-fold receptor density differences expected on a two-dimensional plane, mimicking cell surface density (bottom). C) Scatterplots of pseudotyped virus infectivity as a function of calibrated abundance scores of ACE2 variants. The semi-translucent red, green, and blue lines denote LOESS fits to the points. The scatterplots in the bottom row overlay these LOESS curves. The horizontal purple line denotes the infectivity of template plasmid, which encodes a null ACE2 sequence lacking its start codon. D) Schematic demonstrating the STIM1 sequence variants, their nucleotide identifiers, and their anticipated cell toxicity phenotypes (top), as well as a key denoting how STIM1 abundance level is expected to correspond to cell toxicity (bottom). E) Histograms showing the distribution of cell survival scores, separated by STIM1 variant. F) Overlaid LOESS curves fitting data-points denoting cell survival as a function of STIM1 calibrated abundance score.

To test this concept, we tagged the WT ACE2 Kozak library construct with A, while the known loss-of-binding ACE2 variant D355N was tagged with C, and a minor infection-enhancing variant I21N was tagged with G (**Fig 7B**). The ensuing cells were pooled, infected with one of three pseudotyped viruses, selected with hygromycin, sequenced, and computationally separated for analysis using the nucleotide identifier. Like earlier, we observed no WT ACE2-dependent enhancement to infection by VSV-G pseudotyped virus, but dose-dependent enhancement to infection for viruses coated with SARS-CoV or SARS-CoV-2 spike (**Fig 7C, red**). D355N ACE2 cells in the same well were poorly infectable, although they showed a slight uptick to infectivity at the highest abundance levels, suggesting that high overexpression can partially functionally compensate for poor binding (**Fig 7C, blue**). In contrast, I21N ACE2 cells exhibited enhanced infectivity relative to the other variants, particularly for SARS-CoV spike (**Fig 7C, green**).

We next applied this strategy to STIM1, which is involved in sensing depletion of endoplasmic reticulum calcium to conformationally rearrange and trigger influx of cytoplasmic calcium from the extracellular environment. We recently demonstrated that the myopathy-causing gain-of-function R304W variant, and the immunodeficiency-associated R429C variant, exhibit contrasting effects on cell-growth and survival when overexpressed in HEK 293T landing pad cells (Kamath & Matreyek, 2025). Kozak libraries for these STIM1 variants were tagged by invariant nucleotide identifiers and expressed in HEK 293T landing pad cells (**Fig 7D**). DNA was amplified prior to and following recombination and growth in culture, sequenced with high throughput DNA sequencing, and analyzed for relative cell survival over time. Overall, cells expressing R304W depleted in culture relative to WT, while cells with the R429C variant became enriched (**Fig 7E**).

Separation of the variants by relative expression levels revealed dose-dependent contributions to variant effect. R304W caused overt toxicity of this variant regardless of the level it was expressed. Increasing dosage of WT STIM1 protein abundance caused reduced survival, indicative of cellular dysregulation occurring through gross overexpression (**Fig 7F**). Despite its overall high level of cell survival, R429C yielded a potentially complex curve of dose-based effect. Our results demonstrate that simultaneous modulation of protein sequence and steady-state abundance are a potentially valuable approach to understanding the impacts of protein coding variants, either as a quantitative assay itself or as an unbiased means for assay optimization.

## Discussion

In this work, we provide improved measurement of the changes to steady-state protein abundance possible through modulation of the Kozak sequence. This information improves retrospective observational analysis, as it aids in functional interpretation of existing genetic variation. We furthermore demonstrate that synthetic variation of the Kozak sequence with nucleotide libraries can be leveraged to tie differences in protein abundance with functional effect, a relationship not captured in most cell-based assays.

The wealth of additional existing information about the Kozak sequence helps to infer the generality of our results. We observed high correlation between our data and the existing gold-standard (Noderer *et al*, 2014). As this dataset was collected in murine cells, this suggests that our results will likely apply across mammals. Our data also correlated, albeit to a reduced extent, with a smaller dataset collected in insect cells (Acevedo *et al*, 2018).

We observed improved correlation with the gold-standard dataset when values imputed from a model trained on our calibrated abundance scores were used. This may be attributable to more than one factor. Our sort-seq workflow exhibits experimental variability, error from which will likely impact individual experimental measurements more than individual predictions from a computational model based on that data. Furthermore, neither computational model likely accounts for sequences with higher-order outlier effects, such as “GCGCGCATG”, which was predicted by both models to cause higher steady-state protein abundance than experimentally determined. To limit any associated concerns, we chose to perform all subsequent analyses with our calibrated abundance scores.

We demonstrate the utility of the Kozak scores for data interpretation by identifying clinically observed Kozak sequence germline variants predicted to reduce steady-state protein abundance greater than 4-fold, in genes known to potentially cause disease when the ensuing protein function is reduced. There are a handful of examples of Kozak sequence changes being associated with human disease, such as a −6G>C change in β-globin exacerbating thalassemia presentation in an Italian family when present as a compound heterozygous with other thalassemia promoting alleles (De Angioletti *et al*, 2004), and a different −6G>C change in the Kozak sequence of the transcription factor GATA4 which was associated with atrial septal defects in a Dutch family (Mohan *et al*, 2014).

While we looked specifically at predicted expression-reducing Kozak germline variants in clinically actionable genes, expression-enhancing Kozak variants may also dysregulate physiology in certain genes. Somatic Kozak variants may also be examined, such as the tumor-associated Kozak variants that may be contributing to oncogenesis (Noderer *et al*, 2014). By capturing how Kozak sequence relates to protein-steady state abundance, our dataset may aid future development of more intricate models relating genomic sequence to functional effect, such as expression quantitative trait loci. This information may also aid therapeutic reversion of disease-associated Kozak variants through base-or prime-editing of endogenous loci (Ambrosini *et al*, 2022; Xie *et al*, 2023).

Cells naturally regulate functional output through programmed changes to protein synthesis, such as during differentiation (Kristensen *et al*, 2013). We leveraged this overall strategy of modulating protein synthesis to enact functional changes at the cellular level, albeit through targeted perturbation of the rate of translation initiation. This method is experimentally convenient since it only requires programming of a small stretch of nucleotides, easily introduced with an oligonucleotide primer during PCR. Regulation of protein translation initiation is the final step of protein expression, and is thus tightly linked to protein function. This likely contributes to our observation that cell-to-cell variability in protein expression was drastically reduced with programmed control of translation initiation, in contrast to the drastically increased variability observed with transcriptional control of the doxycycline-inducible promoter.

Our calibrated abundance scores established in this work also provide reference values for future multiplexed genetic assays measuring the relationship between protein abundance and cellular effect. The concept of reactant modulation is also well established for *in vitro* biochemical experiments, which often leverage titration of purified components, but we have lacked the tools to perform similar modulation in cell based experiments. This is particularly important for proteins that exhibit membrane localization and/or multimerization, as such situations often involve binding avidity which is typically difficult to re-create authentically with isolated components *in vitro*. The binding interactions underlying virus entry can be an example of this, where cell-based assays with viral particles can reveal functional interactions between virus and host membrane glycoproteins that are not observed when tested *in vitro* as monomeric subunits (Shukla *et al*, 2021). Despite the D355N variant abrogating binding or infection with all previously tested assay formats (Shukla *et al*, 2021; Ren *et al*, 2022; Bhattacharjee *et al*, 2021), we still observed a slight uptick to SARS-CoV-2 spike pseudotyped virus infection conferred by ACE2 D355N expressed at the highest levels, suggesting that binding avidity is still counteracting the drastic loss of binding affinity conferred by this mutant.

Affinofile cells, which use small molecules as inducers to modulate the cell surface expression of CXCR4 and CCR5, have been a valuable tool for HIV research (Johnston *et al*, 2009), and our system furthers these capabilities. With Kozak-determined translation regulation being genetically encoded, it allows for multiplex genetic approaches not possible through chemical induction. Transcriptional regulation used in the Affinofile system likely mimics the characteristics of our own doxycycline titration experiments, where inducer titration modulates mean protein expression at the population level while exhibiting much higher single-cell heterogeneity than our translational regulation system. Since our experiments were performed with HEK293T landing pad cells purposefully engineered to accept diverse cDNA cargos, additional protein interactions could easily be tested by targeted genomic integration of the relevant coding sequences, requiring vastly less cell engineering than other non-modular methods.

Our work highlights how the same functional assay can differ depending on whether each sequence is being tested in isolation as a homogenous population or as a pool multiplex experiment. Lentiviral particles coated with spike glycoprotein and entering host cells via interactions with ACE2 exhibited a maximum limit to the dose-dependent increase to infection when each sample was tested individually as homogenous cultures, while this limit was not encountered when the assay was performed as a pool library. For the many real-life cases where phenotypic variation is occurring in an otherwise mixed pool of cells, it is possible that the multiplexed format will capture information in a manner more relevant to the underlying biology under investigation than the traditional arrayed format.

Our results point to the vast potential of simultaneously modulating protein sequence and abundance level when performing the cell based functional assays. We could have tested the three variants of ACE2 and STIM1 as separate Kozak libraries, but the fully-pooled format possible by using the −6 Kozak position as the coding variant identifier allowed us to mitigate batch effects and similar sources of experimental error. Even larger sequence-abundance multiplex assays of variant effect will necessitate additional barcoding capacity through larger nucleotide identifier sections, but readily achievable through existing molecular methods.

Regulation of protein translation initiation is a convenient and effective method for functional regulation, both in natural and synthetic contexts. Our work provides calibrated, quantitative values that improve interpretation of naturally occurring variants that may contribute to disease. Our experiments also establish the methodological framework for genetic control of protein translation as a molecular rheostat, wherein control of signal intensity can be combined with libraries of protein coding variants to enable understanding of the combinatorial intersection of protein sequence and abundance to cellular function.

## Materials and Methods

### Cell culture

The generation of the HEK 293T LLP-Int-IRES-iCasp9-Blast Bxb1 landing pad cell line used in this work was previously described (Shukla *et al*, 2021). Cells were cultured in D10 medium (Dulbecco’s modified Eagle’s medium supplemented with 10% fetal bovine serum, 100 U/mL penicillin, and 0.1 mg/mL streptomycin). To induce transcription from the landing pad promoter, the cells were switched to D10X medium (D10 supplemented with 2 μg/mL doxycycline). Long-term passaging of unrecombined landing pad cells was performed with D10X medium supplemented with 20 μg/mL blasticidin, to select against cells that silence the landing pad locus. All cell detachment steps were performed with a 0.25% solution of Trypsin-EDTA (Corning, 25-053-CI).

### Recombinant DNA construction

Recombinant plasmid DNAs were generated using Gibson Assembly. DNA fragments were amplified from 40 ng of plasmid DNA using 0.33 μM each of forward and reverse primers and 2X Kapa HIFI HotStart ReadyMix. The PCR conditions included an initial denaturation at 95°C for 5 minutes, followed by nine cycles of 98°C for 20 seconds, 65°C for 15 seconds, and 72°C for 8 minutes, concluding with a final extension at 72°C for 5 minutes. The resulting amplicons underwent DpnI digestion with 20 U of enzyme at 37°C for 2 hours to remove template DNA. Reactions were purified using a Zymo clean and concentrator kit, and 2 μL of the eluted product was combined with 2 μL of GeneArt Gibson Assembly enzyme mix (Thermo Fisher, A46629). The mixture was incubated at 50°C for 30 minutes before transformation into chemically competent *E. coli* DH10β cells, which were subsequently plated on LB-agar supplemented with 150 μg/mL ampicillin and incubated overnight. Colonies were screened by overnight growth in LB media containing 150 μg/mL ampicillin, followed by plasmid isolation using a GeneJet Plasmid Miniprep kit (Thermo Fisher, K0503). The presence of the desired transgene was verified in each candidate clone through Sanger sequencing, with whole plasmid sequencing performed on all selected clonal variants utilized in the study.

To generate the ACE2 Kozak libraries, Gibson assembly reactions were prepared in 30 μL volumes consisting of 40 ng of template DNA, 0.33 μM of the designated primer pair, 10 μL of nuclease-free water, and 15 μL of Kapa HiFi HotStart ReadyMix polymerase. For each library, four PCR reactions were prepared. The assembled plasmid products were pooled and purified with a Zymo clean and concentrator kit. 5 μL of the cleaned product was combined with 5 μL of GeneArt Gibson assembly master mix and incubated at 50°C for 60 minutes. The assembled plasmid product was purified with a Zymo clean and concentrator kit and eluted in 10 μL of nuclease-free water. Cleaned PCR products were electroporated into *E. coli* DH10β cells (New England BioLabs, C3019I). Each bacterial pellet corresponding to a library was resuspended in 1 mL of sterile LB medium, and appropriate volumes (8, 16, 32, and 64 μL) were plated on LB-agar plates containing ampicillin to assess colony-forming units. The remaining bacterial culture was used to inoculate 200 mL of LB with 150 μg/mL ampicillin, followed by plasmid purification using a NucleoSpin Plasmid kit (Macherey-Nagel, NC0389395). To confirm successful construction, 6-8 bacterial colonies from LB plates were picked randomly, purified using the GeneJET Miniprep kit, and sequence-verified by Sanger sequencing using an Applied Biosystems 3730 Genetic Analyzer.

The G868A_AttB_[mutATG]ACE2-miRFP670_IRES_mCherry-H2A-P2A-PuroR plasmid served as the template for the standard ACE2 Kozak library containing 4096 variants using primers KAM3549 and KAM3665, while the same plasmid was used to create mini libraries with primers KAM4516 and KAM3549. For generating ACE2 Kozak mutant libraries, the following plasmids and primer pairs were utilized: the I21N mutant library was created using the G868A_AttB_[mutATG]ACE2-I21N_A-miRFP670_IRES_mCherry-H2A-P2A-PuroR plasmid with primers KAM4519 and KAM3549; the K31D mutant library was constructed using the G868A_AttB_[mutATG]ACE2-K31D_B-miRFP670_IRES_mCherry-H2A-P2A-PuroR plasmid with primers KAM4517 and KAM3549; and the D355N mutant library was generated from the G868A_AttB_[mutATG]ACE2-D355N_C-miRFP670_IRES_mCherry-H2A-P2A-PuroR plasmid using primers KAM4518 and KAM3549. Primer sequences used for creating the sequence variant libraries can be found in **Supplementary Table 1**.

The following plasmids previously deposited in Addgene were used in this work: pNK005C_BattB_mkozak_STIM1 _IRES_mCherry-H2A-P2A-PuroR (Addgene plasmid #200639) (Roelle *et al*, 2023), G1088E_pLenti-CMV-mNeonGreen-2A-HygroR (Addgene plasmid #216279) (Shukla *et al*, 2024). pMD2.G was a gift from Didier Trono (Addgene plasmid # 12259; http://n2t.net/addgene:12259; RRID:Addgene_12259). The SARS-CoV and SARS-CoV spike expression vectors were previously described (Shukla *et al*, 2021). Additionally, the following plasmids were deposited in Addgene as part of this work: G790A_AttB_ACE2-miRFP670_IRES_mCherry-H2A-P2A-PuroR (Addgene plasmid # 237440) and G868A_AttB_[mutATG]ACE2-miRFP670_IRES_mCherry-H2A-P2A-PuroR (Addgene plasmid # 237441).

### Genomic integration of attB plasmids into landing pad cells

For the recombination of clonal ACE2-encoding attB plasmid constructs, the aforementioned HEK 293T Bxb1 landing pad cells were transfected in 6-well plates. 600,000 cells were plated into each well, and transfected with a mixture of 1.2 μg plasmid of Bxb1 attB-encoding plasmid and 5 μL of Fugene 6 transfection reagent (Promega, E2692), in 100 μL total of Opti-MEM (Gibco, 31985070). Three days after transfection, non-recombined landing pad cells were negatively selected with the addition of 10 nM AP1903 (ApexBio, B4168) to activate iCasp9. Non-silenced, recombined cells were subsequently positively selected with the addition of 1 μg/mL puromycin (InvivoGen, ANTPR1). Successfully selected cells were maintained in D10X media with 1 μg/mL puromycin to prevent transgene silencing during culture.

The miRFP670 Kozak library was recombined and sorted in four separate instances to create the four replicates analyzed in our dataset. For the first biological replicate, 120,000 HEK 293T cells were plated per well in a 24-well plate, and each well was transfected with a mixture of 2 μg ACE2 Kozak library plasmid and 8 μL Fugene transfection reagent (Promega, E2692), diluted in 60 μL Opti-MEM (Gibco, 31985070). Eight wells were transfected per attempt and subsequently pooled. Flow cytometry analysis prior to selection showed 2.5% mCherry-positive cells, suggesting that approximately 24,000 cells were independently recombined, corresponding to an estimated 6 independent events per Kozak sequence variant. Following positive and negative selection, sequencing revealed that some key Kozak sequences necessary for calibrating our results with our panel of individual variants were missing.

To address the coverage limitations observed in the first replicate, a modified approach was taken for the second replicate. For this replicate, 1 million cells were seeded in a 6-well plate one day before transfection. Cells were transfected with a mixture containing 4 μg ACE2 Kozak library plasmid, 30 ng G790A plasmid, and 1.2 μL Xfect reagent, following the manufacturer’s instructions. After positive and negative selection, approximately 8,500 cells from each of the following clonal plasmid-expressing cell lines, G1100A [TTCATCATG], G1101F [GACGACATG], G1102A [CGTCCAATG], G1103A [GCGCGCATG], G1104B [TGGTCAATG], and G1106A [TTGCACATG], were mixed with 10 million library-expressing cells to achieve a diverse range of protein abundance. For the third biological replicate, 1 million cells were seeded in a 6-well plate one day before transfection. Cells were transfected with a mixture containing 4 μg ACE2 Kozak library plasmid, 30 ng G790A plasmid and 30 ng of each Kozak sequence plasmids (G1100A, G1101F, G1102A, G1103A, G1104B, and G1106A), and 1.2 μL Xfect reagent, following the manufacturer’s protocol. Following recombination, cells were subjected to positive and negative selection as described before. This approach was designed to enhance the diversity of protein abundance linked to Kozak sequence variations. For both of these replicates, ~ 170,000 cells were collected during each 4-way sort and the cells were cultured for ~48hrs before their genomic DNAs were extracted.

The fourth recombination experiment used an independent miRFP670 Kozak library, where miRFP670 was C-terminally fused to the hygromycin resistance gene. Here, cells were transfected in two separate wells of a 6-well plate using Xfect reagent. Each well containing 1 million cells was transfected with 4 μg of Hygromycin Kozak library plasmid mixed with 1.2 μL Xfect reagent.Following recombination, cells underwent positive and negative selection as mentioned above. Roughly 130,000 cells were collected in each bin during 4 way sorting and were cultured for ~48hrs before collecting them for gDNA isolation.

Additionally, each ACE2 coding variant-associated Kozak mini library was individually transfected into two separate wells of a 6-well plate using the Xfect transfection reagent (Takara Bio, 631318). Each well containing 400,000 cells was transfected with 4 μg of mini-library plasmid, mixed with 1.2 μL of Xfect reagent, following the manufacturer’s protocol. The cells for each mini-library transfection were pooled prior to selection, and flow cytometry showed roughly 5% of the cells to express mCherry. Assuming each library contained 1,024 Kozak sequence variants, each Kozak mini-library is estimated to have yielded 39 independent recombination events, on average, per Kozak sequence. The library was then negatively and positively selected as described above. The separate mini-library recombinations were pooled prior to infection with pseudotyped virus.

### ACE2 cell surface immunofluorescence staining

Cells expressing the original ACE2 Kozak library were trypsinized to obtain a single-cell suspension. The cell suspensions were washed with P5 buffer (PBS + 5% FBS). Cells were incubated with 1:50 dilution of ACE2 primary antibody (Abcam, ab272500) for 45 minutes at 4°C, followed by two washes with P5 buffer. Subsequently, cells were incubated with a 1:300 dilution of goat anti-rabbit IgG Alexa Fluor 488 secondary antibody (Invitrogen, A32731) for 40 minutes at 4°C in the dark. After two final washes with P5 buffer, the cells were resuspended in 200 μL of P5 buffer, and analyzed by flow cytometry.

### Analytical flow cytometry

Cells were detached from the plates using trypsin, resuspended in P5 buffer, and analyzed by flow cytometry with a Thermo Fisher Attune NxT cytometer. mTagBFP2 was excited with a 405 nm laser, and emitted light was collected after passing through a 440/50 nm band pass filter. EGFP was excited with a 488 nm laser, and emitted light was collected after passing through a 530/30nm band pass filter. mCherry was excited with a 561 nm laser, and emitted light was collected after passing through a 620/15 nm band pass filter. iRFP670 and miRFP670 were excited with a 638 nm laser, and emitted light was collected after passing through a 720/30 nm band pass filter.

### Cell sorting by iRFP670 fluorescence

Cells recombined with the Kozak variant libraries were sorted into 4 separate bins of gradation for near-infrared fluorescence intensity using a BD FACS ARIA SORP (BD Biosciences). mTagBFP2 was excited with a 405 nm laser, and emitted light was collected after passing through a 450/50 nm band pass filter. EGFP was excited with a 488 nm laser, and emitted light was collected after passing through a B515/20 nm band pass filter. mCherry was excited with a 561 nm laser, and emitted light was collected after passing through a 610/20 nm band pass filter. miRFP670 was excited with a 640 nm laser, and emitted light was collected after passing through a 670/30 nm band pass filter.

The pooled cell library was detached from the culture plates using Trypsin-EDTA, and subsequently transferred into D10 media. Following centrifugation at 300 x g for 3 min, the supernatants were discarded, and cell pellets were resuspended in 1 ml of cold P5 buffer. Live cells were gated using the area values of forward and side scatter. Singlet cells were subsequently gated using the forward scatter height and area values. Recombined cells were subsequently gated by subsetting for cells with high mCherry and low mTagBFP2 fluorescence. A histogram of cell count based on iRFP670 fluorescence intensity was generated, upon which 4 gates were drawn to separate the population into quartiles. Approximately 120,000 cells were collected per bin, for 5 total replicates.

### Pseudotyped virus infection assays

Pseudotyped virus particles were produced in otherwise unmodified HEK 293T cells seeded into 6-well plates. Each well plated with 1 million cells was transfected with a mixture containing 1 μg of PsPax2 (Addgene #12260), 1 μg of the lentiviral transfer vector G1088E_pLenti-CMV-mNeonGreen-2A-HygroR, and 1 μg of viral envelope plasmid (VSV-G, SARS-CoV spike, or SARS-CoV-2 spike), and PEI-Max MW 40,000 (PolySciences, CAS Number: 49553-93-7). Twelve hours after transfection, the medium was removed and the cells were washed with 1X PBS before incubating the cells with 2 mL D10 media. The viral supernatant was collected each day over the next 72 hours. For each pseudotyped virus, supernatants from nine wells of a 6-well plate were pooled. Upon collection, the media was centrifuged at 300 x g for 3 minutes, and the clarified supernatant was transferred to a fresh tube. Different volumes of clarified supernatant were used for infection: 8 mL and 24 mL for SARS1, 8 mL and 32 mL for SARS2, and 2 mL and 8 mL for VSVG. These were mixed with 6 million library cells seeded in 100 mm dishes. Four days post-infection, cells were examined under a fluorescence microscope to assess mNeonGreen expression. Subsequently, 10% of the infected library cells were harvested for genomic DNA (gDNA) isolation, while the remaining 90% were subjected to hygromycin selection (185 μg / mL and 400 μg / mL concentrations). These concentrations are much higher than normally necessarily for 293T cells, and was likely inflated due to a combination of high culture cell densities during selection, along with the presence of a large proportion of hygromycin-resistant cells within the cell population, which likely metabolized standard hygromycin concentrations, thus protecting sensitive cells at standard hygromycin concentrations (50 μg / mL). After hygromycin selection, the remaining cells were trypsinized, pelleted, and frozen for subsequent genomic DNA extraction.

### ACE2 amplicon generation and high throughput sequencing

Genomic DNA was isolated from each sample using the JetFlex™ Genomic DNA purification kit (Thermo Fisher, A30700). A section of DNA spanning the recombination junction, and including the Kozak sequence preceding the ACE2 open reading frame, was amplified from the genomic DNA extracts using PCR. For each 50 µL PCR reaction, the following components were used: 2 µg genomic DNA (quantified by nanodrop), 0.25 µM forward primer containing i7 index barcode and 0.25 µM reverse primer containing i5 index barcode, and 25 µL of Phusion plus high fidelity PCR mix (Thermo Fisher, F548L). A total of five PCR reactions were performed per condition, which each tube initially incubated at 95 °C for 3 minutes, followed by 25 cycles of 95 °C for 15 seconds, 60 °C for 15 seconds, 72 °C for 30 seconds, followed by a final 72 °C incubation for 1 minute. PCR products were mixed with 6x DNA loading dye (NEB, B7024S) and separated on 1.5% agarose TAE gel. Bands corresponding to the expected 293 bp amplicon size were excised and extracted using the Freeze and Squeeze kit according to the manufacturer’s instructions (Bio-Rad, 7326166). Eluted PCR products were purified using a Zymo clean and concentrator kit (Zymo Research, D4003). Amplicon concentrations were measured using Qubit dsDNA BR assay kit (Life technologies, Q32853). Amplicons from all the experimental conditions were pooled in equimolar ratios and sequenced on an Illumina NextSeq550 with a 75-cycle high output kit. Oligonucleotide sequences for PCR and custom Illumina primers for sequencing are listed in **Supplementary Table 1**.

### STIM1 Kozak library generation and cell survival assay

The STIM1 Kozak library was generated using an oligonucleotide primer (Integrated DNA technologies) where the −6 to −1 nucleotides preceding the STIM1 start codon was encoded by a degenerate XNNNNN segment, where X is the invariant nucleotide at the −6 position for barcoding (A for WT, C for R429C, and G for R304W) (**Fig 7A, 7D**). Gibson cloning, electroporation, and individual library purification was performed as described for ACE2, above. The three plasmid libraries were individually prepared, subsequently mixed in equimolar ratios, and transformed into DH10β *E.coli*, which was midiprepped to yield the final library introduced into the landing pad cells.

For each replicate, two 100 mm plates, each with 3.5 million HEK 293T landing pad cells, were transfected with the plasmid library using the Xfect transfection reagent, as described above. Cells were harvested 5, 7, and 9 days following transfection. Genomic DNA was extracted from each time point using DNeasy Blood and Tissue Kit (Qiagen, 69506), and quantitated with a Qubit fluorometer using the Qubit Broad Range dsDNA DNA quantification kit (Thermo Fisher, Q32853).

Both the midiprepped plasmid library, as well as the genomic DNA from each time-point, were amplified by PCR to create the material necessary for high throughput DNA sequencing. The reaction conditions were similar to that of ACE2, except 200 ng of genomic DNA (as quantified by the broad range dsDNA Qubit kit) was used as the starting material. Amplicons were separated on a 1.5% agarose TAE gel, excised, extracted, and quantitated by broad range dsDNA Qubit as described above. The vast majority of samples were sequenced on an Illumina NextSeq 550 using a Mid Output 150 cycle kit by the CWRU Genomics Core. The first three replicates of the STIM1 Kozak library were sequenced using the Illumina MiSeq, generated through submission to the Amplicon-EZ service offered by Genewiz / Azenta. Oligonucleotide sequences for PCR and custom Illumina primers for sequencing are listed in **Supplementary Table 1**.

### Identification of clinical germline Kozak variants

The current list of reference sequence identifiers for GRCh38 was downloaded from NCBI. The reference RNA sequences were translated and cross referenced with the reference protein sequences for each gene to identify the first exact protein match, and the 6 nucleotide positions preceding the start codon in the RNA were returned as the Kozak sequence. As a secondary approach, the longest possible translated protein sequence was identified for each RNA, and if a methionine was present, the 6 nucleotides preceding the first methionine were used to identify the Kozak sequence.

The March 2025 release of ClinVar was used to identify observed clinical variants that occur within the Kozak sequence. The data was isolated to the GRCh38 assembly to match the reference sequences, and entries were filtered for variants between position −6 and position −1. Kozak sequence variants that overlapped with introns were identified by the script, but not used for subsequent analysis due to possible additional complications of transcript splicing. Kozak variant sequences were then paired with the associated RNA sequence using the NM accession. The variant was then checked against the Kozak generated by the first method and applied if a positional match was found, and both the original and new sequence were returned. If the first method generating the Kozak returned no sequence, or if the positional mutation did not match the generated Kozak, then the Kozak returned by the second method was used instead, and the variant was returned using that approach instead. Variants involving microsatellites were ignored in this analysis. 1,826 explicit Kozak variants were identified from the 2,558 total variants within the Kozak region identified in the ClinVar database (**Supplementary Table 2**).

## Data analysis

For short-read sequencing generated by the Illumina NextSeq, the forward and reverse reads were paired using PEAR (Zhang *et al*, 2014). Kozak sequences were counted from the paired reads using Enrich2 (Rubin *et al*, 2017), and filtered to exclude reads possessing nucleotides with quality scores less than 20. For longer-read sequencing generated by the Illumina MiSeq, generated through submission to the Amplicon-EZ service offered by Genewiz / Azenta, the reads were paired as above, except the paired reads were subsequently processed with a custom Python script “Extract_Kozak.py”, which identified true Kozak library reads from template molecule reads, and for each Kozak read, extracted the Kozak sequence found in that read. All subsequent analyses of the sequencing outputs were performed in R version 4.3.1 (2023-06-16). An R markdown file with code to recreate the analysis (Kozak.Rmd), along with the relevant data files and the various python scripts for the initial processing of FASTQ files, can be found in the MatreyekLab Github repository under “MatreyekLab/Kozak”. A table of the fastq files generated and analyzed for this work is included as **Supplementary Table 3**.

The Kozak sort-seq scores were calculated similarly to previous work (Matreyek *et al*, 2018, 2021), with some minor changes. For samples sequenced to sufficient read depth, a density-plot of sequence counts normally yields a bimodal distribution, with one peak stemming from a log-normal distribution of counts with a median of 10 or greater, largely reflecting sequences of physical molecules amplified and sequenced from the sample, and a separate peak with a count of 1, largely reflecting sequences derived from sequencing error. We harnessed this property to computationally remove erroneous sequence counts. For each bin, we identified the local minima between these two peaks, and removed sequence counts below this sample-specific threshold value, to yield a dataset of filtered counts.

For each sequenced sample, the filtered count of each variant was divided by the sum of counts recorded in that sample to obtain variant-specific frequencies (F_v,bin_). For replicates 1 and 2, which both included separately sequenced amplification “technical” replicates derived from the same sorted genomic DNA, the technical replicate frequencies were averaged to yield an average frequency for each variant, which was used as the F_v,bin_ value. These frequencies were used to calculate a weighted average for each variant for each replicate (W_v,rep_), using the following formula.

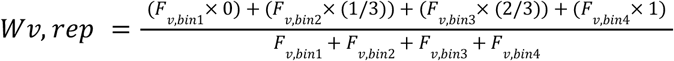

Thus, all weighted average values ranged from a value of 0 to 1. Pairwise comparisons of variant-specific weighted averages are shown in **Supplementary Fig 2**. Once the weighted average values were calculated for each variant for each replicate, they were log_10_ transformed, averaged, and the base-10 exponential calculated to yield the geometric mean weighted average value. Correspondingly, the standard deviation of the log_10_ transformed values were calculated and used to calculate the upper and lower 95% confidence intervals, which were later transformed as a base-10 exponential to accompany the geometric mean weighted averages.

To calibrate these weighted averages to MFI equivalent values, we generated cells recombined with ACE2-miRFP670 constructs encoding 16 different Kozak sequences, which were measured for miRFP670 fluorescence intensity using analytical flow cytometry. A linear model was first generated to establish the conversion factor enabling scaling of our weighted average scores to MFI-equivalent values. While accurate for values between 1% and 100% of the consensus Kozak, the linear model performed poorly for extremely high or low values, with the minimum and maximum scaled values being 0.0003 and 3.89; both of these values were outside of the range physically observed with this assay. We thus used logistic regression to approximate the linear model, but furthermore bound all scaled values between 0.004 and 1.34. These resulting values are referred to as the calibrated scores. These scores are provided in **Supplementary Table 4**.

Enrichment values for infection experiments were initially filtered similarly to the sort-seq scores, wherein we identified the local minima between the two peaks observed in a density plot representation of sequence counts, and removed sequence counts below his sample-specific threshold value, to yield a dataset of filtered counts. The counts for each variant were divided by the total counts for that sequence sample, to yield sample-specific variant frequencies. For infection samples that included separately sequenced amplification “technical” replicates derived from the same sorted genomic DNA, the geometric means of the technical replicate frequencies were used. Each hygromycin-selected infected sample had a corresponding unselected sample. For each pair, we divided variant-specific frequencies following selection by those prior to selection, obtaining an enrichment ratio. The STIM1 survival scores were calculated similarly, with the exception that the ratio was the quotient of each Kozak variant frequency recombined into cells divided by the frequency of that variant in the sequenced plasmid library. The scores for these functional assays are found in **Supplementary Tables 4 and 5**.

## Acknowledgements

The authors wish to thank Simone Edelheit and Milena Zelembaba of the Genomics Core Facility of the CWRU School of Medicine’s Genetics and Genome Sciences Department. The Cytometry & Imaging Microscopy Shared Resource of the Case Comprehensive Cancer Center was supported by NIH grants P30CA043703 and S10OD021559. Nidhi Shukla thanks the Flora Stone Mather Center for Women at CWRU for providing a research and professional development grant to support the presentation of this work at the Discover BMB 2023 conference.

## Data Availability

The authors affirm that all data necessary for confirming the conclusions of the article are present within the article, figures and tables. Key plasmids have been deposited to and available via Addgene. Code to recreate analysis is available on Github (https://github.com/MatreyekLab/Kozak). Raw FASTQ data can be found at the NCBI GEO repository under accession number GSE295498.

## Funding

Funder Grant reference Number Author

National Institutes of Health GM142886 Kenneth A Matreyek

National Institutes of Health AI178151 Kenneth A Matreyek

National Institutes of Health AI161275 Anna M Bruchez

The funders had no role in study design, data collection and interpretation, or the decision to submit the work for publication.

## Author contributions

Nidhi Shukla: Investigation, Methodology, Writing - review and editing Nisha D Kamath: Investigation, Methodology, Writing - review and editing John C Snell: Investigation, Data curation, Formal analysis, Methodology, Writing - review and editing Anna M Bruchez: Writing - review and editing Kenneth A Matreyek: Conceptualization, Methodology, Investigation, Data curation, Formal analysis, Visualization, Writing - review and editing, Supervision, Funding acquisition, Methodology, Project administration.

