## Supplementary Fig 1 for "Kozak sequence libraries for characterizing transgenes across expression levels"

### Supplementary Tables

**Supplementary Table 1.** List of primers and associated sequences for library generation, library amplification, and subsequent high throughput sequencing.

**Supplementary Table 2.** Table of Kozak variant ClinVar entries and their predicted impacts to protein expression.

**Supplementary Table 3.** List of high-throughput sequencing fastq files deposited and analyzed in this work.

**Supplementary Table 4.** Table of sort-seq weighted averages, calibrated scores, random-forest imputed scores, and enrichment values following selection for cells infected with pseudotyped lentivector particles. Individual infection values and assessed mean-fluorescent intensities of control constructs are incorporated into the table.

**Supplementary Table 5.** Table of infection values for Kozak libraries of WT, I21N, and D355N ACE2, when mixed with VSV-G, SARS-CoV spike, and SARS-CoV-2 spike pseudotyped lentivector particles.

**Supplementary Figures**

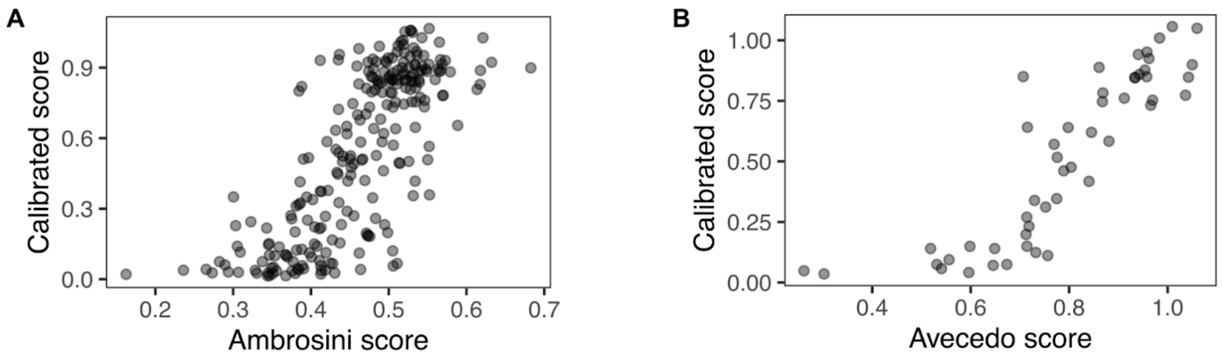


**Supplementary Fig 1**. Comparison of calibrated scores with scores calculated in diverse model systems. Comparison with A) scores calculated by Ambrosini *et al*, with lentivirally transduced HEK 293T cells, and B) scores calculated by Avecedo *et al*, tested in Drosophila Kc167 cells.

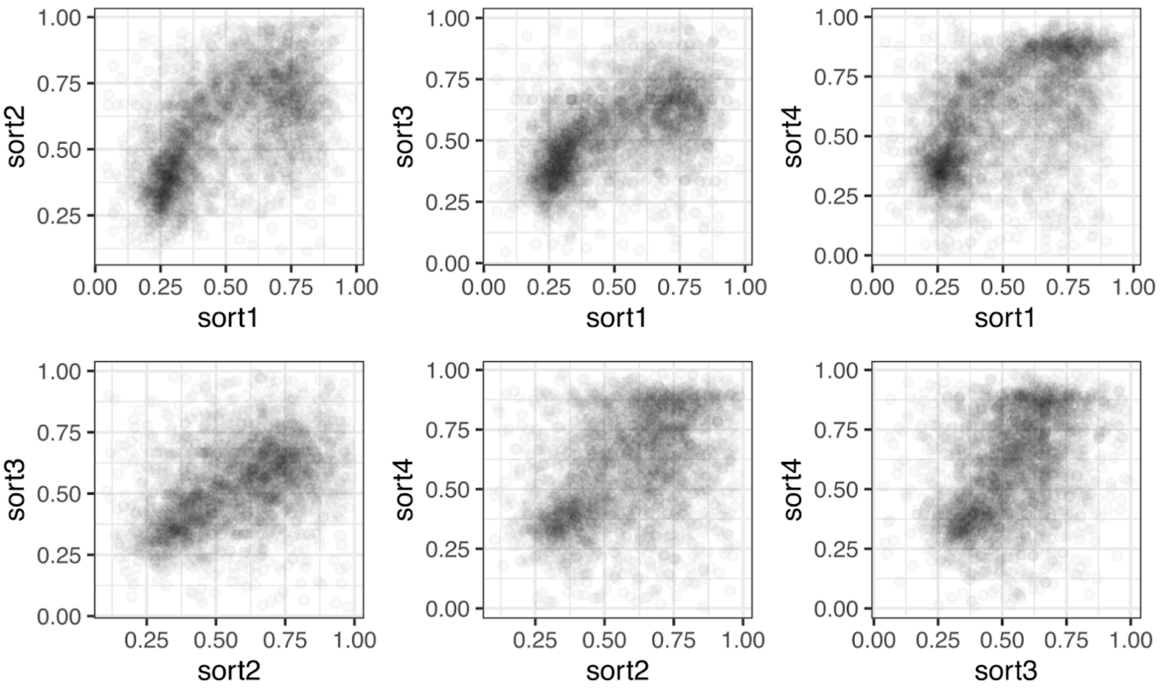


**Supplementary Fig 2**. Pairwise comparison of variant-specific weighted average values for each sort-seq replicate.
